# Synanthropic flora in the vicinity of the mediaeval Castle Kolno, Stare Kolnie near Brzeg, southwestern Poland

**DOI:** 10.1101/2022.12.26.521972

**Authors:** Romuald Kosina, Lech Marek

**Affiliations:** Faculty of Biological Sciences, University of Wrocław, Przybyszewskiego 63, 51-148 Wrocław, Poland; Institute of Archaeology, University of Wroclaw, Szewska 48, 50-137 Wroclaw, Poland

**Keywords:** Mediaeval castle, cabbage, garden-field cultivation, weeds, ruderals

## Abstract

The mediaeval castle Kolno situated near the village Stare Kolnie served as a custom house at the confluence of two rivers, Budkowiczanka and Stobrawa. Numerous diaspores of plants were obtained from the archaeological excavations, from the layers of the 14th-15th centuries A.D. The excavations were located near the former access road, where increased human activity has affected the composition of the fossil macroremains of plants. Two species of cabbage cultivated in small fields near the castle were recognised. Diaspores of weeds and ruderal plants were deposited at the site. The most frequent were: *Solanum nigrum, Setaria pumila, Chenopodium album, Rumex acetosella, Polygonum lapathifolium* and *Urtica dioica*. The botanical set of fossil diaspores was composed of plant species associated with anthropogenic habitats and showing the dispersion dynamics in various micro-niches within them.

## 1. Introduction

The analysis of archaeobotanical data on foods consumed in the Middle Ages covers the countries of western and northern Europe, where the study of fossil macroremains has progressed to a more advanced stage (Greig 1983). These data indicate that cabbage was the predominantly cultivated vegetable. This finding was confirmed by later archaeobotanical analyses (Greig 1986; Wiethold 1995; Speleers & van der Valk 2017) as well as by comparative data of fossil materials from the literature of the era (Helweg 2020). Macrofossils from eastern Europe also confirmed the cultivation of this plant (Wasylikowa 1978; Latałowa et al. 1998; Kosina & Marek 2021; Badura et al. 2015;). The cultivation of cabbage and other vegetables was accompanied by weeds, whose habitat range also included ruderal places with loose and fertile soils. Thus, assigning or not assigning weeds identified in the fossil layers of diaspores to vegetable crops depends on whether they are obligatory or optional in relation to the cultivated plant.

In Poland, after World War II, the southern areas, including Lower Silesia, were a particularly good area for cabbage cultivation. Cultivation of this vegetable can be limited to simple agrotechnical treatments, which does not reduce its yields (Chroboczek 1966). Probably this practice has also been known and important for a mediaeval farmer. The modern map (Fig. 1) shows the higher elevations in the vicinity of the castle Kolno, where cultivation was possible also in the Middle Ages. The water network of this area was very rich, and the dispersion of plants from typical riparian communities into the vicinity of human habitats was highly probable. This is confirmed by fossil remains (Kosina unpubl.).

**Fig. 1.**
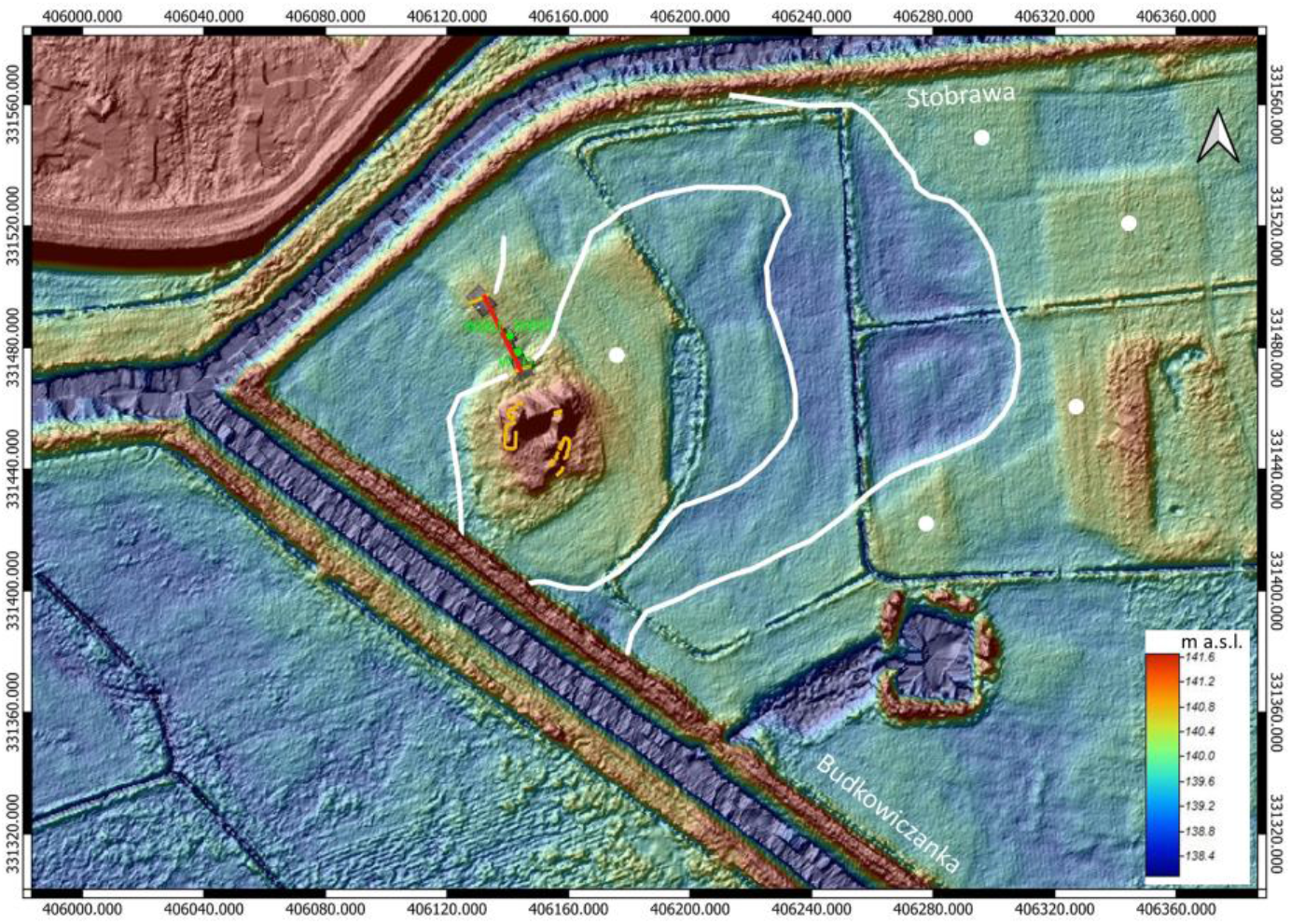
Digital elevation model based on LIDAR, castle Kolno. Marked with a white line is the probable original outline of the river meander and point bar on which the castle is located. Highlighted in red is the orientation and location of the bridge leading from the castle to the village. Marked with yellow lines are castle walls and the original banks of the castle’s moat documented in archaeological trenches. Marked with green is the position of the analysed samples. Higher elevations where crop cultivation was possible are marked by white dots. By L. Marek

Mediaeval weed infestation of gardens or vegetable fields could be related to the contemporary state; however, scarce data are available on this subject. Anioł-Kwiatkowska (1974) and Siciński (2000) studied individual fields of head cabbage and indicated a high frequency of *Chenopodium album* in the so-called secondary weed infestation. This infestation of weeds occurs in the second half of the vegetation period, probably because of the minimal anti-weed effect of currently used herbicides. New information on the dynamics of the synanthropic flora, useful for comparison with fossil materials, should be obtained from the junction of garden and field crops with anthropogenic habitats (Marshall & Moonen 2002; Piórek & Krechowski 2010).

The present study aimed to support the data presented earlier (Kosina & Marek 2021) based on new archaeobotanical analyses. This involved the cultivation of cabbage in gardens or small fields and weed infestation of these crops in contact with ruderal habitats. The research material was the sediment in the moat; thus, it was largely mixed, unlike the stored crops from the fields, which are mainly homogeneous (Kosina 1977; Lityńska-Zając 2018). This fact limits the syntaxonomic analyses of the macroremains obtained from the moats.

## 2. Materials and methods

### 2.1. Archaeology

Castle Kolno is located south of the village of Stare Kolnie, Opole district, inside the river fork of Budkowiczanka distributary and the Stobrawa River (the coordinates of the castle’s center according to the PUWG1992 coordinate system: x: 406148,5; y: 331450,2). The geomorphology of the site is characterised by the presence of alluvial sand- and gravel - deposits forming the terraces of the Stobrawa- and Budkowiczanka-Rivers. Holocene river fans comprising organic and loamy silts and sandy loam soil, typical for floodplains were recorded on the site. The castle is located on a point bar composed of alluvium deposits accumulated by the river, which was altered afterward by anthropogenic, alluvial, and aeolian processes. The fortress was most probably raised already in the 13th century as testified by the oldest archaeological evidence collected on the site. Originally located on the border of the Duchies of Brzeg (since 1419 the Duchy of Legnica and Brzeg) and Opole the fortress played a very important economic, administrative and military role. It guarded a custom house where charges were levied on forest goods transported down the river, such as wax, timber, honey, etc. The castle was besieged and destroyed on July 13th, 1443, and the following days during the so-called succession war in Silesia (Ermisch 1876, p. 61; for further reading on the history of the castle please see Marek 2014, pp. 131-132). As confirmed by the archeological evidence collected to date, the castle was most probably never rebuilt nor inhabited after this event. The most destructive for the castle’s ruins was stone quarrying and recycling in the 18th century which led to the dismantling of still-existent castle walls to a considerable extent. Therefore the reconstruction of the stone structure and its original layout, even after several archaeological campaigns on the site remains a challenge. The methods involved during our investigations apart from regular excavations with the use of floatation procedures and sifting screens was the analysis of spatial data recorded with the use of Leica Total Station device – 407 Modell and GPS RTK device: Hi-Target V30 GNSS equipped with Q-Mini controller and Hi-Target Hi-RTK Road program, 3D scanning, as well as archaeological surface- and geomagnetic surveys. Essential for the mapping of finds was the interpretation of spatial data produced by the LIDAR scanning.

Frequent river flooding of the area led to the obliteration of elevated terrain features containing anthropogenic archaeological strata. The latter destroyed and heavily distorted were moved by the water to form the fill of the original terrain depressions. The excavations in the area of the castle’s moat in 2012 revealed, that under these distorted archaeological deposits there is an original strata sequence. On top of it, there is a thick river-silt deposit resulting from the drying out of the moat after the destruction of the fortress in 1443. No archaeological evidence of settlement on the site was documented in this layer. Underneath there was the original deposit of organic matter, and a rich archaeological record documenting the everyday activities, character, and social status of the inhabitants and people visiting the fortress. All of it had great significance for the dating of these layers. Valuable in this context is the ring tree-dated construction of the bridge leading from the castle to the north, where the village is located (Fig. 1 and 2). Several bridge posts found in the archaeological trenches date to the beginning of the 14th century, and the branches that fell into the moat to ca 1325. According to dendrochronology, the castle’s moat was used from the 1st quarter of the 14th century onwards until the beginning of the 1440s. Of course, a castle’s moat is far from an orderly stratigraphic sequence where alluvial processes and objects falling into it disrupted original deposits. It was possible however to observe that the lowest layers contained the oldest archaeological evidence dated to the beginning of the 14th century such as the earliest coins (see Marek & Paszkiewicz 2012), a seal stamp, and early stove tiles with images of the first recorded owner of the castle Boleslaw III the Generous (Marek 2017, pp. 73-75, Figs. 1 and 4). The youngest layers forming the original moat fill contained several small coins dated to the 20s and early 40s of the 15th century and arms and armour finds related to the siege of 1443. Among the latter, there were numerous crossbow bolt heads and entirely preserved bolts, fragments of mediaeval firearms and leaden fire-arm projectiles, elements of mediaeval brigandine armour, sword fragments, and daggers (see Marek 2020). After several excavation seasons focusing on the moat area, we may conclude, that archaeological evidence correlates stunningly well with the results of ring tree dating. The original water level in the moat at its highest is estimated to be 138.9 m a.s.l. based on the recorded sediments. The samples analysed in an already published study (Kosina & Marek 2021) were taken from ceramic vessels of early 15th-century chronology, whereas the ones being the subject of this paper are from a 14th-century jug found in the lowest layer of the moat fill. Additionally, a soil sample was taken from the same trench in which the vessel was recorded (Table 1, a moat deposit).

**Table 1.**
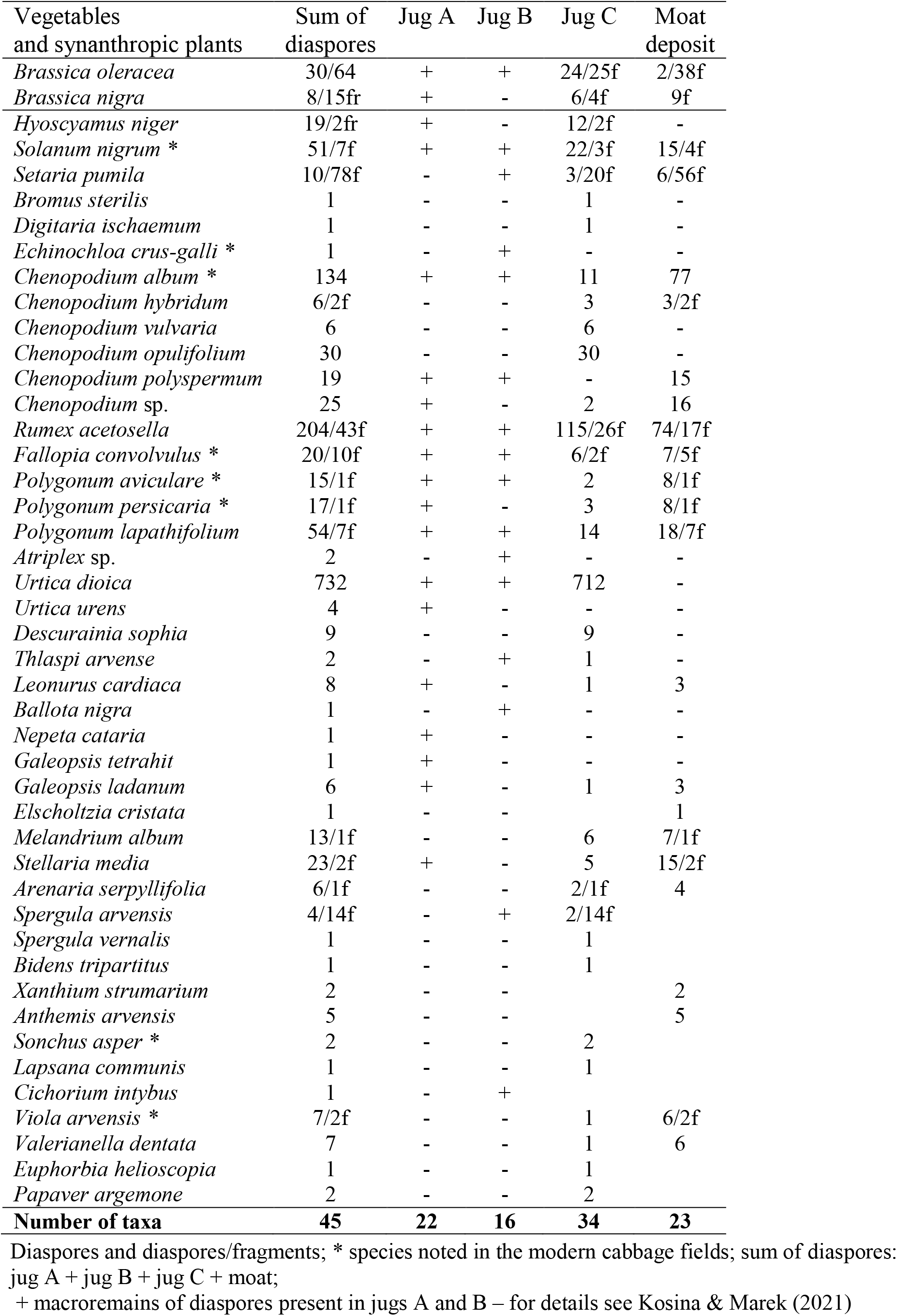
Fossil diaspores of crops, weeds and ruderals from the Castle Kolno

| Vegetables<br>and synanthropic plants | Sum of<br>diaspores | Jug A | Jug B | Jug C | Moat<br>deposit |
| --- | --- | --- | --- | --- | --- |
| *Brassica oleracea* | 30/64 | + | + | 24/25f | 2/38f |
| *Brassica nigra* | 8/15fr | + | - | 6/4f | 9f |
| *Hyoscyamus niger* | 19/2fr | + | - | 12/2f | - |
| *Solanum nigrum* * | 51/7f | + | + | 22/3f | 15/4f |
| *Setaria pumila* | 10/78f | - | + | 3/20f | 6/56f |
| *Bromus sterilis* | 1 | - | - | 1 | - |
| *Digitaria ischaemum* | 1 | - | - | 1 | - |
| *Echinochloa crus-galli* * | 1 | - | + | - | - |
| *Chenopodium album* * | 134 | + | + | 11 | 77 |
| *Chenopodium hybridum* | 6/2f | - | - | 3 | 3/2f |
| *Chenopodium vulvaria* | 6 | - | - | 6 | - |
| *Chenopodium opulifolium* | 30 | - | - | 30 | - |
| *Chenopodium polyspermum* | 19 | + | + | - | 15 |
| *Chenopodium* sp. | 25 | + | - | 2 | 16 |
| *Rumex acetosella* | 204/43f | + | + | 115/26f | 74/17f |
| *Fallopia convolvulus* * | 20/10f | + | + | 6/2f | 7/5f |
| *Polygonum aviculare* * | 15/1f | + | + | 2 | 8/1f |
| *Polygonum persicaria* * | 17/1f | + | - | 3 | 8/1f |
| *Polygonum lapathifolium* | 54/7f | + | + | 14 | 18/7f |
| *Atriplex* sp. | 2 | - | + | - | - |
| *Urtica dioica* | 732 | + | + | 712 | - |
| *Urtica urens* | 4 | + | - | - | - |
| *Descurainia sophia* | 9 | - | - | 9 | - |
| *Thlaspi arvense* | 2 | - | + | 1 | - |
| *Leonurus cardiaca* | 8 | + | - | 1 | 3 |
| *Ballota nigra* | 1 | - | + | - | - |
| *Nepeta cataria* | 1 | + | - | - | - |
| *Galeopsis tetrahit* | 1 | + | - | - | - |
| *Galeopsis ladanum* | 6 | + | - | 1 | 3 |
| *Elscholtzia cristata* | 1 | - | - |  | 1 |
| *Melandrium album* | 13/1f | - | - | 6 | 7/1f |
| *Stellaria media* | 23/2f | + | - | 5 | 15/2f |
| *Arenaria serpyllifolia* | 6/1f | - | - | 2/1f | 4 |
| *Spergula arvensis* | 4/14f | - | + | 2/14f |  |
| *Spergula vernalis* | 1 | - | - | 1 |  |
| *Bidens tripartitus* | 1 | - | - | 1 |  |
| *Xanthium strumarium* | 2 | - | - |  | 2 |
| *Anthemis arvensis* | 5 | - | - |  | 5 |
| *Sonchus asper* * | 2 | - | - | 2 |  |
| *Lapsana communis* | 1 | - | - | 1 |  |
| *Cichorium intybus* | 1 | - | + |  |  |
| *Viola arvensis* * | 7/2f | - | - | 1 | 6/2f |
| *Valerianella dentata* | 7 | - | - | 1 | 6 |
| *Euphorbia helioscopia* | 1 | - | - | 1 |  |
| *Papaver argemone* | 2 | - | - | 2 |  |
| **Number of taxa** | **45** | **22** | **16** | **34** | **23** |

Diaspores and diaspores/fragments; \* species noted in the modern cabbage fields; sum of diaspores: jug A + jug B + jug C + moat; + macroremains of diaspores present in jugs A and B – for details see Kosina & Marek (2021)

**Fig. 2.**
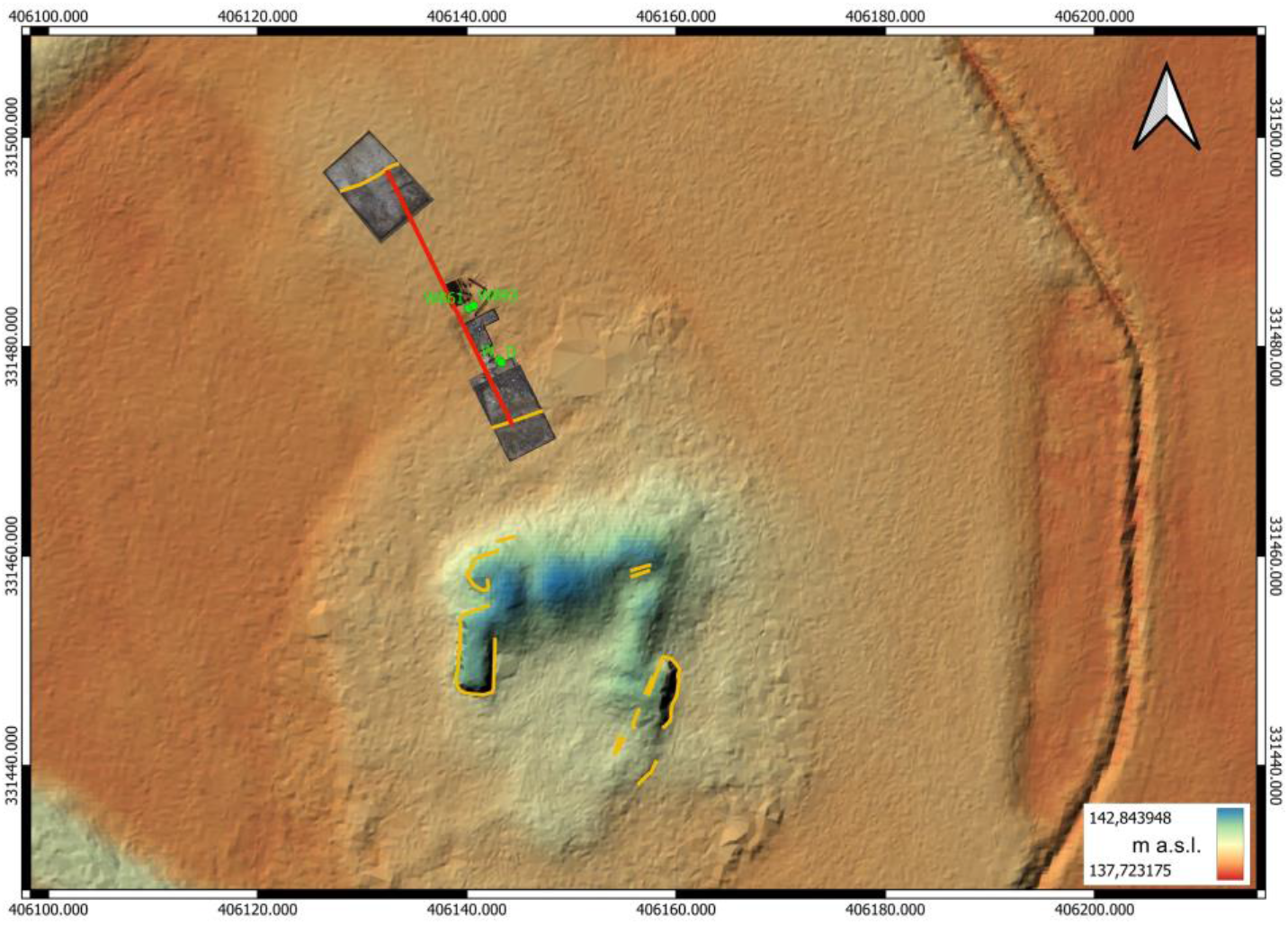
Digital elevation model based on LIDAR, castle Kolno. Highlighted in red is the orientation and location of the bridge leading from the castle to the village. Marked with yellow lines are castle walls and the original banks of the castle’s moat documented in archaeological trenches. Marked with green is the position of the analysed samples. By L. Marek

### 2.2. Archaeobotany

The fossil plant material was collected during archaeological excavations from the former moat surrounding the castle (Fig. 1, 2). Diaspores were selected on several sieves from fossil layers in the moat and from three jugs: A, B and C. Diaspores from vessels A and B were described earlier (Kosina & Marek 2021), and they were used to summarise the data in Table 1, without showing their numbers. Diaspores were analysed based on several carpological keys and a rich collection (RK) of contemporary and fossil diaspores.

## 3. Results and Discussion

The summarised data are presented in Table 1. The detection of new diaspores (Jug C and moat deposit) confirmed the presence of seeds of two cabbage species in the fossil material. On the modern map of the studied area (Fig. 1) there are marked areas of higher elevations, where mediaeval cultivation was possible. These areas corresponded to small fields or garden plots. Thus, it is a strong basis to conclude that some agricultural activity with the cultivation of *Brassica oleracea* and *B. nigra* occurred in this area.

Cabbage cultivation has an ancient tradition in Europe. This is reflected in the work De agri cultura written by Marcus Porcius Cato, in which *B. oleracea* was considered the best vegetable (Hammer et al. 2018). The introduction of *B. oleracea* to England also dates back to the time of the Roman invasion of the island. Historical records indicate its widespread cultivation, consumption and medicinal use (Mitchell 1976). In mediaeval Poland, seeds of both species, more often *B. nigra*, were recorded at many archaeological sites from the Middle Ages, in the period concurrent with the functioning of castle Kolno. Thus, *B. oleracea* was frequently consumed in mediaeval Gdańsk in the 15th century (Badura et al. 2015). Many *B. nigra* seeds were also served from Elbląg from the 13th to 14th century (Latałowa et al. 1998). However, cabbage seeds were not numerous on Wawel Hill in Cracow (Wasylikowa 1978). Very large amounts of cabbage seeds, namely *B. nigra* and *B. rapa*, have been identified in strata 13th-15th c in Brussels (Speleers & van der Valk 2017) and Ferrara, northern Italy, from the 14th century to the end of the 15th century A.D. (Bandini Mazzanti et al. 2005). Similarly, in Kiel, numerous diaspores of *B. nigra* were found in the material collected from the moat (Wiethold 1995). In contrast, cabbage seeds were not identified among the numerous crop diaspores identified in the Prague 14th c. moat layers (Beneš et al. 2002). These findings indicate that, despite differences in some sites (Prague), cabbage cultivation in mediaeval Europe was common and confirms Greig’s earlier analysis of northern Europe (Greig 1983). This set of mediaeval cabbage cultivation sites should also include the outskirts of castle Kolno. It is assumed that in the outskirts of the castle, cabbages were grown on small plots on the sandy sediments of the Stobrawa and Budkowiczanka rivers. These poor soils were probably enriched with fertile silts during spring floods or supplemented with manure. Cabbage was sown after the flood waters receded.

Cabbage cultivation can be simplified and limited to soil harrowing. During sowing in early spring and in the longer period of lower temperature (approximately5°C), the plants do not form a head and show premature flowering in the first year of cultivation. Contemporary data indicate that cabbage is not sensitive to weed infestation (Chroboczek 1966). These requirements of the plant facilitated its cultivation in the vicinity of the castle, moreover, premature flowering (bolting) yielded seeds identified in the fossil layers. Table 1 includes a large set of diaspores of synanthropic plants, weeds and ruderal plants. They constitute only a part of the taxa identified at the Kolno site and are related mainly to the cultivation of root and vegetable crops and other anthropogenic habitats in contact with them. Diaspores from the two previously analysed vessels, jugs A and B (Kosina & Marek 2021), only are summarised in Table 1. The plants with 10 diaspores or more were *Hyoscyamus niger, Solanum nigrum, Setaria pumila, Ch. album, Ch. opulifolium, Ch. polyspermum, Rumex acetosella, Fallopia* (*Polygonum*) *convolvulus, Polygonum aviculare, P. persicaria, P. lapathifolium* (*nodosum*), *Urtica dioica, Melandrium album* and *Stellaria media*.

Among diaspores of root and garden crop weeds and ruderal plants obtained from the mediaeval layers on the Wawel Hill in Cracow, the most numerous were diaspores, which were also frequent in the castle Kolno. These are: *Setaria pumila, Echinochloa crus-galli, Fallopia convolvulus, Chenopodium album, Melandrium album, Polygonum persicaria, Polygonum aviculare, Urtica dioica, Solanum nigrum* and *Hyoscyamus niger* (Wasylikowa 1978). Quantitative differences between the Wawel and Kolno locations resulted from the difference in terms of their size and the volume of the examined fossil samples. However, the repetition of certain plants in the form of numerous diaspores at various archaeological sites is evident. It is also confirmed by data from mediaeval sites in Denmark (Jensen 1986). The set of diaspores from the mediaeval moat (1200-1550 A.D.) in Svendborg (Jensen 1979) reflects the diversity of plants identified in the Kolno moat. *Urtica dioica* and *Solanum nigrum* were the most numerous. Plants with the most numerous diaspores could be treated as those of local origin (Latałowa et al. 1998).

Few studies have analysed the weed infestation of modern cabbage crops in Poland. Anioł-Kwiatkowska (1974) presented extensive data on the synanthropic flora from the northwestern part of Lower Silesia. Among the weeds in the field of head cabbage (*B. oleracea* var. *capitata*), the author lists the following species also found in the fossil material from Kolno: *Sonchus asper, Ch. album, Sinapis arvensis, Centaurea cyanus, Convolvulus arvensis, P. persicaria, F. convolvulus* and *Viola arvensis*. On the other hand, in the cultivation of head cabbage in the vicinity of Łęczyca in the *Echinochloo*-*Setarietum* association, Siciński (2000) noted the following species: *Echinochloa crus*-*galli, P. lapathifolium, C. cyanus, V. arvensis, Ch. album, Capsella bursa*-*pastoris, S. nigrum*, and *P. aviculare*. In both cases, the fields were heavily infested with *Galinsoga parviflora*. Table 1 shows the weeds of cabbage crops reported by Anioł-Kwiatkowska and Siciński. References to modern fields, however, are of limited value for comparison with fossil materials; this is because modern weed flora is greatly degraded by chemical treatments. Unfortunately, data of fields whose agricultural status can be compared to old fields are unique. An example is Tymrakiewicz’s study on weeds in the agricultural fields of Lower Silesia before the period of herbicide use. This study, however, does not provide any information on cabbage cultivation (Tymrakiewicz 1952). Lityńska-Zając (2005) indicated that weeds of root crops were less than those of cereal crops in mediaeval fossils. The author lists *Euphorbia helioscopia, S. nigrum*, and *P. persicaria* among the former group. The author also considered that modern syntaxonomy has limited application to fossil materials. Because we are investigating a mixture of diaspores (Kolno) for fossil accumulation in the moats, the use of syntaxonomic description here would be a methodological error. In contrast, a stored set of single-species diaspores with a rich set of weeds, which are non-mixed and dated for a narrow time interval, can be characterised by the above method (Kosina 1977).

Among the species listed in Table 1, many of them can be assigned to ruderal habitats. These include *H. niger, S. nigrum, Ch. album, Urtica urens, Descurainia sophia, Thlaspi arvense, Leonurus cardiaca, Ballota nigra, Nepeta cataria, Stellaria media* and *Xanthium strumarium*. The latter species in the area of the castle was probably associated with riparian vegetation. However, it may have preferred ruderal sites, as indicated by comparative analyses of plant variability in natural and ruderal habitats (Blais & Lechowicz 1989). Significant amounts of its diaspores, probably used for dyeing fabrics, were discovered at a site in Switzerland dated for the early mediaeval period (Brombacher & Hecker 2015).

Valuable materials for comparison with the ruderal flora of earlier periods can be obtained from places where traditional farming methods are likely to be used. An example of this may be the flora of ruderal habitats in abandoned villages in the area of the Kampinos National Park (Kirpluk 2011). Several ruderal archaeophytes were found in this area: *Ballota nigra, C. bursa*-*pastoris, D. sophia, Urtica urens, V. arvensis, E. crus*-*galli, L. cardiaca, C. cyanus, F. convolvulus* and *S. nigrum. C. bursa*-*pastoris* and *D. sophia* were the most common species. Some of these species are listed in Table 1. Species such as *Ch. album, P. persicaria, P. aviculare, M. album* and *R. acetosella* occurring in the vicinity of the castle were noted with a high frequency at numerous archaeological sites (Trzcińska-Tacik & Wasylikowa 1982). The habitat range of ruderal plants, apart from various sites strongly disturbed by humans, also includes arable fields. They can grow in various vegetable crops, including cabbage and root crops. A dynamic exchange of diaspores occurs between both types of habitats: ruderal and cultivated. For example, dispersion of the ruderal *D. sophia* into contemporary agricultural fields in Lublin Province, eastern Poland, was particularly frequent in winter oilseed rape crops (Kapeluszny &, Haliniarz 2010).

The distribution of plants in various environmental mosaics, namely arable fields, fallow land, orchards, gardens, and communication routes, first occurs at the junctions of these environmental puzzles. The importance of these junctions, e.g. roadsides, was analysed by Piórek & Krechowski (2010). The authors found that presently, the weeds of root crops, such as *C. bursa*-*pastoris, S. media, Anchusa arvensis, Ch. album, E. crus*-*galli* and *S. pumila*, most often penetrate the roadside. This dispersion of diaspores certainly existed in the Middle Ages near the castle Kolno, where different, but small in area, crops could be adjacent to each other. The castle Kolno was linked to the neighbouring Stare Kolnie village (Alt Köln) by a road running across the bridge over the moat in the north direction (Fig. 1, 2). There are archaeological excavations in this section near the hill of the castle; consequently, it is normal to find the presence of ruderal plants in the excavations near the activity of people on the road and in the castle. Marshall & Moonen (2002) emphasise the fact that a specific micro-mosaic of plants forms on the vegetation belts at the junctions, which may be a refugium of flora from earlier periods. From such microhabitats (field margins), the species penetrate into fields, thus becoming weeds or inhabiting ruderal places. The inhabitants of the outer settlement and the castle had to address this dynamic environmental micro-mosaic situation. Lososová et al. (2006) analysed the characteristics differentiating field communities from ruderal ones and indicated that the ruderal environment is more variable than the field one. Ruderal plants have been classified by the authors as: biennials or perennials, … wind-pollinated, flowering in mid-summer, reproducing both by seeds and vegetatively, dispersed by wind or humans, … species with high demands for light and nutrients. The synanthropic plants listed in Table 1 meet the above-mentioned criteria in many cases.

## 4. Concluding remarks

In the vicinity of the mediaeval castle Kolno dated the 14th-15th century and located in the immediate vicinity of Alt Köln village, there were cultivations of cabbages (*B. oleracea* and *B. nigra*) in the form of small fields or garden plots, probably in a small area at the outskirt of the castle. The identified seeds may originate from the premature flowering of some plants (in Polish ‘pośpiechy’). Among the synanthropic plants from fields and ruderal habitats, e.g. roadsides, the most numerous were *S. nigrum, R. acetosella, P. lapathifolium* (*nodosum*), and *Urtica dioica*. Ruderal habitats also included the following species: *H. niger, U. urens, D. sophia, L. cardiaca, B. nigra* and *X. strumarium*. A dynamic dispersion of diaspores and mutual penetration of taxa certainly occurred between crops and unstable ruderal habitats, which enriched the floristic richness at the junctions.

## Author contribution statement

The section ‘archaeology’ and Figs. 1, 2, were elaborated and written by LM, the ‘archaeobotany’ part and Table 1 were elaborated and written by RK.

## Conflict of interest

The authors declare that they have no conflict of interest.

## Notes

### Competing Interest Statement

The authors have declared no competing interest.

